# The Phagocyte Oxidase Controls Tolerance to *Mycobacterium tuberculosis* infection

**DOI:** 10.1101/232777

**Authors:** Andrew J Olive, Clare M Smith, Michael C Kiritsy, Christopher M Sassetti

## Abstract

Protection from infectious disease relies on two distinct mechanisms. “Antimicrobial resistance” directly inhibits pathogen growth, whereas “infection tolerance” controls tissue damage. A single immune-mediator can differentially contribute to these mechanisms in distinct contexts, confounding our understanding of protection to different pathogens. For example, the NADPH-dependent phagocyte oxidase complex (Phox) produces anti-microbial superoxides and protects from tuberculosis in humans. However, Phox-deficient mice do not display the expected defect in resistance to *M. tuberculosis* leaving the role of this complex unclear. We re-examined the mechanisms by which Phox contributes to protection from TB and found that mice lacking the Cybb subunit of Phox suffered from a specific defect in tolerance, which was due to unregulated Caspase1 activation, IL-1β production, and neutrophil influx into the lung. These studies demonstrate that Phox-derived superoxide protect against TB by promoting tolerance to persistent infection, and highlight a central role for Caspase1 in regulating TB disease progression.

## Introduction

Protective immunity to infectious disease involves functionally overlapping responses that can be divided into two fundamentally different categories (Medzhitov et al., 2012; Schneider and Ayres, 2008). Infection “resistance” refers to functions that directly target the infecting pathogen to prevent its growth and dissemination. Resistance pathways act by a variety mechanisms including disrupting the bacterial niche, serving as metabolic poisons, and sequestering critical nutrients (Hood and Skaar, 2012; Olive and Sassetti, 2016; Pilla-Moffett et al., 2016). In addition, the extent of disease is also influenced by “tolerance” mechanisms that enhance host survival but do not directly impact pathogen growth (Ayres and Schneider, 2008; Jamieson et al., 2013; Weis et al., 2017). Tolerance pathways control a broad range of functions that protect the infected tissues from both the direct cytotoxic properties of the pathogen and inflammation-mediated immunopathology. While it is well appreciated that both resistance and tolerance mechanisms are required to limit disease, the relative importance of these pathways vary for different infections (Medzhitov et al., 2012). Furthermore, since individual immune effectors can promote both tolerance and resistance, the specific role for each mediator can change in different contexts (Jeney et al., 2014; Medzhitov et al., 2012; Meunier et al., 2017; Mishra et al., 2017). In the context of chronic infections, where resistance mechanisms are insufficient and the pathogen persists in the tissue, tolerance is likely to play a particularly important role (Meunier et al., 2017).

Like many other chronic infections, the outcome of an encounter with *Mycobacterium tuberculosis (Mtb)* varies dramatically between individuals (Cadena et al., 2017). Only 5–10% of those that are infected with this pathogen progress to active tuberculosis (TB), and disease progression is influenced by a wide-variety of genetic and environmental factors that could modulate either tolerance or resistance (Chen et al., 2014; Lopez et al., 2003; Tobin et al., 2012). For example, observations from humans and mice indicate that several specific changes in T cell function may contribute to a failure of resistance and disease progression due to loss of antimicrobial resistance (Jayaraman et al., 2016; Larson et al., 2013; Redford et al., 2011). In addition, studies in animal models indicate that a failure of host tolerance, which is necessary to preserve lung function or granuloma structure, influences the extent of disease (Desvignes et al., 2015; Pasipanodya et al., 2010). While these studies suggest that variation in overall tolerance may be an important determinant TB risk, the specific tolerance mechanisms that influence disease progression in natural populations remain ill defined.

During many bacterial infections the production of reactive oxygen species (ROS) by the NADPH phagocyte oxidase (Phox) is essential to protect the host from disease (Segal, 2005). Phox is a multi-protein complex, including the subunits Cybb (gp91) and Ncf1 (p47) that assemble in activated immune cells to produce superoxide radicals by transferring electrons from NADPH to molecular oxygen (Panday et al., 2015). Humans with deleterious mutations in the Phox complex develop a clinical syndrome known as chronic granulomatous disease (CGD). Leukocytes from patients with CGD are unable to kill a number of bacterial pathogens, such as *Staphylococcus aureus* and *Serratia marcescens*, and this defect is associated with the susceptibility to infection with these organisms (Johnston and Baehner, 1970). Because ROS contributes to the microbicidal activity of phagocytes, previous studies in Mtb-infected mice focused on the role of Phox in antimicrobial resistance. However, mice deficient in Phox are able to restrict *Mtb* growth to levels comparable to wild type animals during the initial stages of infection (Deffert et al., 2014a; Jung et al., 2002); (Cooper et al., 2000; Jung et al., 2002). The lack of an obvious antimicrobial role for Phox has been attributed to the expression of mycobacterial defenses, such as the catalase/peroxidase, KatG, which detoxifies ROS directly, or CpsA which prevents Phox localization to the *Mtb* containing vacuole (Colangeli et al., 2009; Koster et al., 2017; Nambi et al., 2015; Ng et al., 2004). These findings have led to the conclusion that ROS produced by Phox are not required for protection to *Mtb* (Nathan and Shiloh, 2000). In contrast, mutations in the *Cybb* gene are strongly associated with susceptibility to mycobacterial disease, including tuberculosis (Bustamante et al., 2011; Deffert et al., 2014a; Khan et al., 2016; Lee et al., 2008). Mutations that specifically reduce Cybb activity in macrophages produce a similar clinical presentation, highlighting the importance of the macrophage-derived ROS in protection from pathogenic mycobacteria (Bustamante et al., 2011). The apparently conflicting data from mice and humans regarding the importance of Phox during *Mtb* infection suggest two possibilities. Either Phox is differentially required to protect against *Mtb* in mice and humans or the ROS produced by Phox is necessary to control immune mechanisms that do not directly modulate bacterial replication.

Here we re-examined the role of Phox in the context of *Mtb* infection. Consistent with previous reports we found that loss of the Phox subunit Cybb does not alter the growth or survival of *Mtb* during infection. Instead, *Cybb*^−/−^ animals suffered from a hyper-inflammatory disease caused by increased activation of the NLRP3-dependent Caspase-1 inflammasome and IL-1-dependent neutrophil accumulation in the lung. Thus, the protective effect of Phox can be attributed to increased tolerance to *Mtb* infection instead of a direct antimicrobial effect. These studies provide a mechanism to explain the association between Phox expression and TB disease in natural populations, and implicate Caspase-1 as an important regulator of infection tolerance.

## Results

### Cybb^−/−^ mice are susceptible to TB disease, but maintain control of bacterial replication

In order to re-examine the role of Phox in mediating protection against *Mtb*, we compared disease progression and the immune responses in wild type and *Cybb*^−/−^ C57BL/6 mice after infection via aerosol with 50–100 bacteria. We found no significant difference in the survival or bacterial levels in the lung between groups of mice up to 3 months following infection, confirming that Cybb is not required for surviving the early stages of *Mtb* infection (Figure 1A). However, after 100 days of infection, *Cybb*^−/−^ infected mice lost an average of 10% of their body weight while wild type animals gained weight (Figure 1B). Histopathological inspection of the lungs also indicated a difference in disease between these groups, with *Cybb*^−/−^ lungs containing larger and less organized lesions than wild type (Figure 1C). These data suggested that wild type and *Cybb*^−/−^ animals might tolerate *Mtb* infection differently even while harboring identical levels of bacteria.

In order to dissect the mechanisms controlling tolerance to *Mtb* disease in *Cybb*^−/−^ mice, we profiled the infected lungs of animals during infection by flow cytometry. We found no significant differences in the numbers of dendritic cells, macrophages, B cells, as well as total T cells between wild type and *Cybb*^−/−^ mice (Figure S1). In contrast, we observed an early and sustained increase of Ly6G^+^ CD11b^+^ neutrophils in the infected lungs of *Cybb*^−/−^ mice (Figure 1D and 1E). A 3–5-fold increase in the total number neutrophils was observed as early as 4 weeks following infection and was maintained throughout the 12-week study.

**Figure 1.**
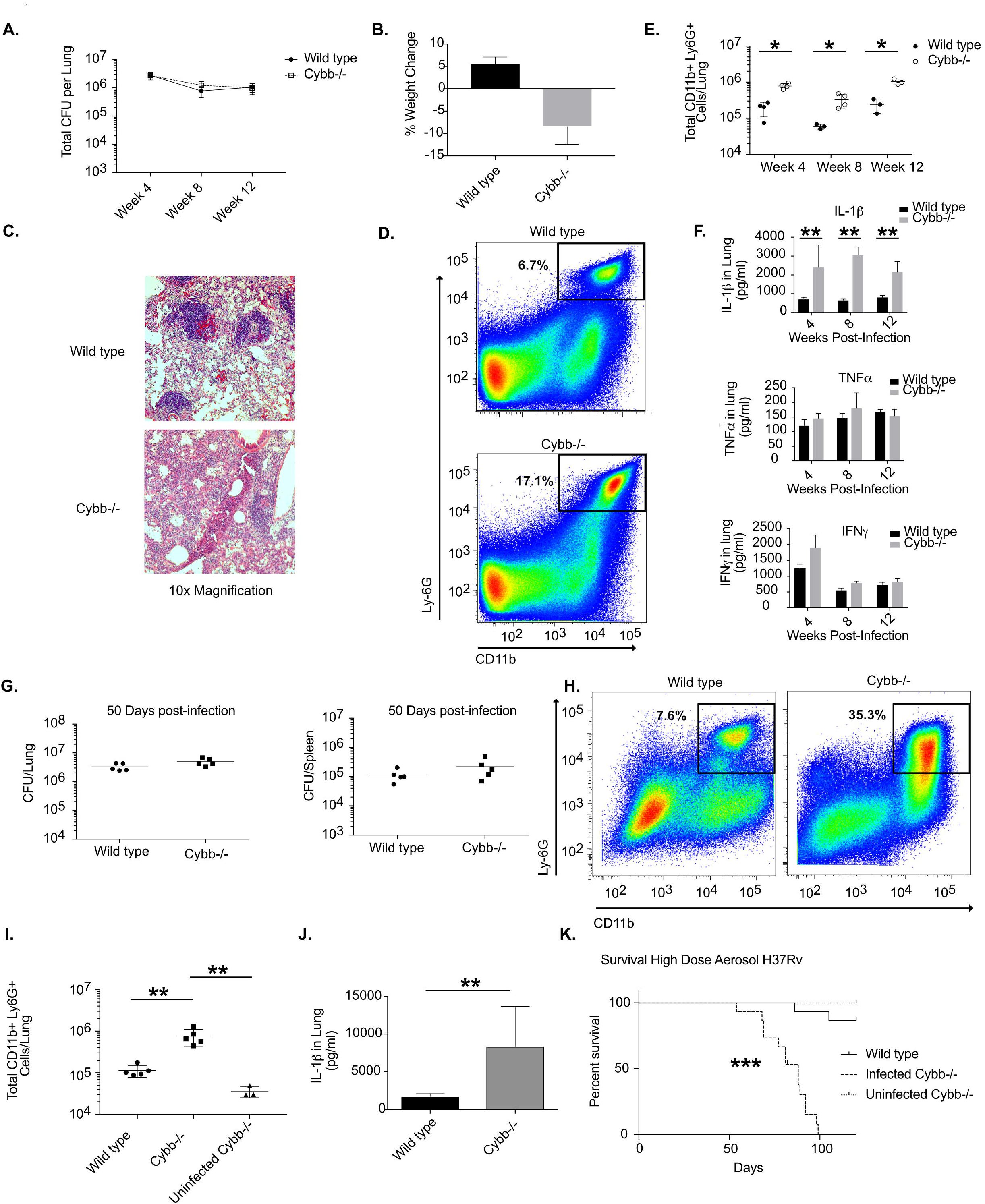
Anti-inflammatory activity of Cybb protects mice from TB disease. **A**. Following low dose aerosol infection (Day 0 of ~50–100 colony forming units, cfu) total bacterial burden (expressed in cfu, mean ^+/−^ s.d.) was determined in the lungs of wild type or *Cybb*^−/−^ mice at the indicated time points. **B**. Percentage weight loss (mean change ^+/−^ s.d) from Day 0 to Day 100 was determined for wild type and *Cybb*^−/−^ mice. **C**. Immunohistochemical staining for Haemotoxylin and Eosin is shown for representative lung sections from wild type and *Cybb*^−/−^ mice at 8 weeks post-infection at 20x magnification. **D**. Representative flow cytometry plot showing increased Ly6G^+^ CD11b^+^ neutrophil recruitment to the lungs of *Cybb*^−/−^ mice 4 weeks following infection (gated on live/singlets/CD45^+^). **E**. Quantification of neutrophil recruitment to the lungs at the indicated time points following infection for wild type and *Cybb*^−/−^ mice is shown as absolute number of Ly6G^+^ CD11b^+^ cells per lung (mean ^+/−^ s.d). * p-value <.05 by unpaired two-tailed t-test. **F**. Lung homogenates from wild type or *Cybb*^−/−^ mice infected for the indicated time were probed for the cytokines IL-1β, IFNγ, and TNFα by ELISA (mean ^+/−^ s.d). Results shown in A-D are representative of 3 independent experiments with 3–5 mice per group. ** p-value <.01 by unpaired two-tailed t-test. **G**. Fifty days following high dose aerosol infection (Day 0 of 5000–7500 CFU) total bacterial burden (expressed in cfu, mean ^+/−^ s.d.) was determined in the lungs and spleen of wild type or *Cybb*^−/−^ mice. **H**. Representative flow cytometry plot showing increased Ly6G^+^ CD11b^+^ neutrophil recruitment to the lungs of *Cybb*^−/−^ mice 50 days following infection (gated on live/singlets/CD45^+^). **I**. Quantification of the absolute number of neutrophils recruited to the lungs 50 days following high dose infection for wild type and *Cybb*^−/−^ mice is shown (mean ^+/−^ s.d). ** p-value <.01 by one-way ANOVA with tukey correction. **J**. Lung homogenates from wild type or *Cybb*^−/−^ mice infected for 50 days following high dose aerosol were probed for IL-1β by ELISA (mean ^+/−^ s.d). ** p-value <.01 by unpaired two-tailed t-test. Results shown in G-J are representative of 2 independent experiments with 5 mice per group. ** p-value <.01 by unpaired two-tailed t-test. **K**. Survival of infected wild type and *Cybb*^−/−^ and uninfected *Cybb*^−/−^ mice was determined following high dose infection. Data are representative of two independent experiments with 1415 mice per group. *** p-value <.001 Mantel-Cox text.

The cytokine IL-1β promotes neutrophil-mediated disease during *Mtb* infection of other susceptible mouse strains (Mishra et al., 2017; Mishra et al., 2013). Similarly, when we assayed cytokine levels in lung homogenates, we noted a dramatic and specific increase in IL-1β concentration in *Cybb*^−/−^ animals compared to wild type at all time points. In contrast, no significant differences were noted for IFNγ or TNFα at any time point between groups. Thus, while the adaptive immune response to *Mtb* appeared to be intact, *Cybb*^−/−^ animals produced excess IL-1β and the concentration of this cytokine correlated with neutrophil infiltration into the lung.

Previous studies have shown that following low dose aerosol infection mice deficient in Phox (both *Cybb*^−/−^ or p47^−/−^) survive for at least 60 days (Cooper et al., 2000; Jung et al., 2002). However, longer infection is likely necessary to determine whether the enhanced disease we noted in *Cybb*^−/−^ mice would result in a survival defect. These long-term survival experiments following low dose aerosol proved difficult, since uninfected *Cybb*^−/−^ mice develop arthritis as they age (Lee et al., 2011). To avoid this confounder, we quantified the survival of mice in a shorter-term study using a high dose aerosol infection. When mice were infected with ~5000 CFU per animal, *Cybb*^−/−^ mice succumbed to disease significantly more rapidly than wild type animals. *Cybb*^−/−^ mice had a median survival time of 88 days while only two out of fifteen wild type mice succumbed during the 120 day study (Figure 1K). In order to distinguish survival effects not related to *Mtb* infection, a cohort of uninfected age-matched *Cybb*^−/−^ mice were maintained for the duration of the experiment. None of these animals required euthanasia over the 120 days and no animals included in this experiment developed arthritis.

During this high-dose study, we also examined a cohort of mice 50 days following infection and found identical levels of bacteria in the lungs and spleen between wild type and *Cybb*^−/−^ groups (Figure 1G). Consistent with our earlier findings, *Cybb*^−/−^ mice showed a significant increase in neutrophils and IL-1β in the lung (Figure 1H–J). We also found minimal levels of IL-1β and neutrophils in uninfected *Cybb*^−/−^ lungs indicating that these phenotypes are dependent on *Mtb* infection. Therefore, the loss of Cybb leads to more severe *Mtb* disease that is associated with increased IL-1β levels and neutrophil recruitment, even though the number of viable *Mtb* in the lung did not appear to be affected.

### Cybb controls tolerance to *Mtb* infection

Our initial results suggested that Cybb protects mice by promoting tolerance to *Mtb* infection rather than directly controlling bacterial replication. However, while viable bacterial numbers were similar in wild type and *Cybb*^−/−^ mice, we could not rule out that the course of disease was altered by subtle changes in the dynamics of bacterial growth and death. To more rigorously address this question, we employed two additional animal models that allowed us to differentiate tolerance and direct antimicrobial resistance *in vivo*.

To more formally exclude the possibility that Cybb alters the intracellular growth of *Mtb* during infection, we used a previously optimized mixed bone marrow chimera approach (Mishra et al., 2017). These experiments normalize potential inflammatory differences between wild type and *Cybb*^−/−^ mice allowing us to specifically quantify differences in bacterial control (Figure 2A). Irradiated wild type mice were reconstituted with a 1:1 mixture of CD45.1+ wild type and CD45.2^+^ *Cybb*^−/−^ or wild type cells. Five weeks following infection, both CD45.1+ and CD45.2+ cells were sorted from the lungs and the levels of *Mtb* in each genotype was determined by plating and the purity of populations was determined by flow cytometry (Figure 2B and 2C and Figure S2). We found that the relative abundance of wild type and *Cybb*^−/−^ cells was similarly maintained throughout infection in both the myeloid and lymphoid compartments, indicating that Cybb does not alter cellular recruitment or survival in a cell-autonomous manner. When *Mtb* was enumerated in sorted cells, we found identical levels of H37Rv in wild type CD45.1^+^ and *Cybb*^−/−^ CD45.2+ populations from the same mouse, similar to the results from mice where both populations were reconstituted with congenically mismatched wild type cells. In contrast, when chimeric mice were infected with a ROS-sensitive *ΔkatG* mutant of *Mtb*, we found higher levels of bacteria in *Cybb*^−/−^ cells compared to wild type cells from the same mouse. These data show that the assay is able to detect the cell-autonomous antimicrobial activity of ROS against a KatG-deficient *Mtb* strain, but Cybb-dependent ROS did not restrict the intracellular replication of wild type *Mtb*.

**Figure 2.**
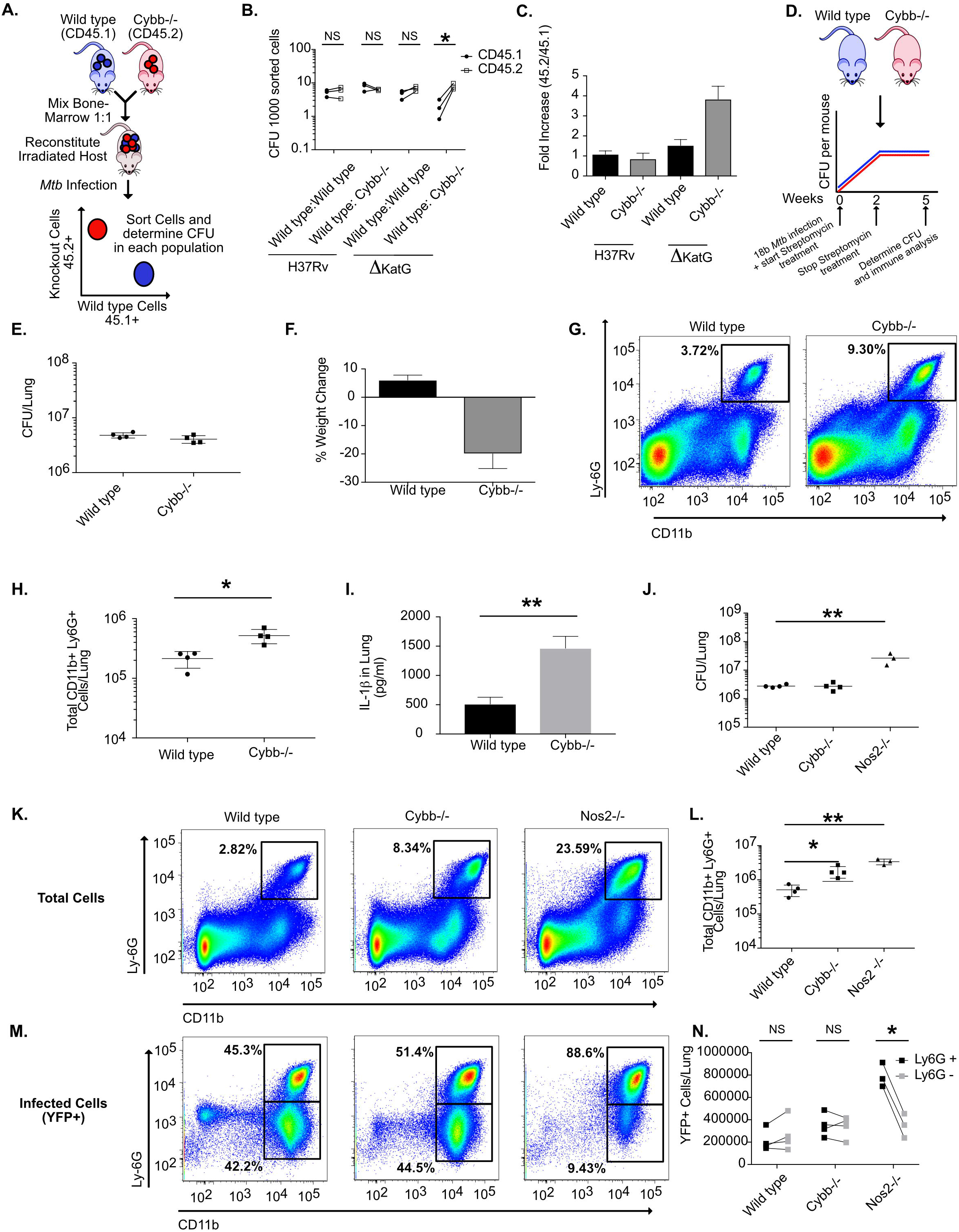
The primary protective role of Cybb is anti-inflammatory. **A**. Schematic for the generation of mixed bone marrow chimeras. Mixed bone marrow chimeras were infected by low dose aerosol with either H37Rv or ΔKatG mutant. Five weeks following infection CFU levels were determined in purified haematopoietic cells of indicated genotypes. **B**. Shown are the normalized CFU per sorted cells in each population from each mouse. * p<.05 by unpaired two-tailed t-test. **C**. The fold-increase of bacterial levels in CD45.2^+^ cells (experimental) compared to CD45.1^+^ cells (wild type control) (mean ^+/−^ s.d.). The results in B and C are representative of three independent experiments with 3–4 mice per group. **D**. Schematic for streptomycin-dependent auxotroph infection. Wild type and *Cybb*^−/−^ mice were infected intratracheally with *Mtb* strain 18b and treated for two weeks daily with streptomycin. Mice were then removed from streptomycin for three weeks halting active growth of the bacteria. **E**. Five weeks after infection the total levels of viable *Mtb* in the lungs was determined by CFU plating on streptomycin (mean ^+/−^ s.d.). **F**. Percentage weight loss (mean change ^+/−^ s.d) from Day 0 to Day 35 was determined for wild type and *Cybb*^−/−^ mice. **G**. Representative flow cytometry plot showing increased Ly6G^+^ CD11b^+^ neutrophil recruitment to the lungs of *Cybb*^−/−^ mice 5 weeks following infection (gated on live/singlets/CD45^+^). **H**. Quantification of neutrophil recruitment to the lungs at the indicated time points following infection for wild type and *Cybb*^−/−^ mice is shown as absolute number of Ly6G^+^ CD11b^+^ cells per lung (mean ^+/−^ s.d). * p-value <.05 by unpaired two-tailed t-test. Data in D-G are representative of 4 independent experiments with 4–5 mice per group. **I**. Lung homogenates from wild type or *Cybb*^−/−^ mice infected with 18b were probed for IL-1β by ELISA (mean ^+/−^ s.d). ** p-value <.01 by unpaired two-tailed t-test. **J**. Following low dose aerosol infection with sfYFP H37Rv (Day 0 of ~50–100 colony forming units, cfu) total bacterial burden (expressed in cfu, mean ^+/−^ s.d.) was determined in the lungs of wild type, *Cybb*^−/−^ or Nos2^−/−^ mice 4 weeks post-infection. ** p-value <.01 by one-way ANOVA with tukey correction. **K**. Shown are representative flow cytometry plots from each genotype of total Ly6G^+^ CD11b^+^ cells in the infected lungs. **L**. Quantification of total neutrophil recruitment to the lungs of the indicated genotypes four weeks following infection (mean ^+/−^ s.d.). ** p-value <.01 * p-value <.05 by one-way ANOVA with tukey correction. **M**. Shown are representative flow cytometry plots from each genotype of infected (YFP^+^ Ly6G^+^ CD11b^+^) cells in the lung. **N**. Enumeration of infected (YFP^+^) neutrophils (Ly6G^+^ CD11b^+^) or monocytes/macrophages (Ly6G-CD11b^+^) in the indicated genotypes. * p-value <.05 by unpaired two-tailed t-test.

To specifically determine if the loss of Cybb decreased tolerance to a given burden of bacteria, wild type and *Cybb*^−/−^ mice were infected with a streptomycin auxotrophic strain of *Mtb* that allows exogenous control of bacterial replication during infection. Streptomycin is provided for the first two weeks of infection, allowing the pathogen to reach the burden observed in a wild type *Mtb* infection. Upon streptomycin withdrawal, the pathogen is unable to replicate but remains viable and able to drive inflammatory responses (Figure 2D) (Honore et al., 1995; Mishra et al., 2017; Mishra et al., 2013). Five weeks after infection, *Cybb*^−/−^ mice lost more weight than wild type animals while harboring identical levels of non-replicating bacteria (Figure 2E and 2F). Lungs from *Cybb*^−/−^ mice contained significantly more neutrophils and higher levels of IL-1β compared to wild type animals (Figure 2G–2I). Thus, even when the need for antimicrobial resistance is obviated by artificially inhibiting bacterial replication, *Cybb*^−/−^ animals continued to exhibit a hyper-inflammatory disease.

The granulocytic inflammation observed in *Cybb*^−/−^ mice was reminiscent of several other susceptible mouse strains. However, the neutrophil recruitment in other models is generally associated with a concomitant increase in bacterial growth (Kimmey et al., 2015; Kramnik et al., 2000; Nandi and Behar, 2011) and a transition of the intracellular *Mtb* burden from macrophages to granulocytes (Mishra et al., 2017). We hypothesized that *Cybb*^−/−^ mice may be able to retain control of *Mtb* replication because the bacteria remain in macrophages. To test this hypothesis, we used a YFP-expressing *Mtb* strain to compare the distribution of cells harboring bacteria in wild type and *Cybb*^−/−^ mice with *Nos2*^−/−^ animals in which *Mtb* replicates to high numbers in association with infiltrating granulocytes (Mishra et al., 2017). Four weeks following infection we found that lungs from both *Cybb*^−/−^ and *Nos2*^−/−^ mice contain higher levels of IL-1β and neutrophils than wild type animals, although the loss of *Nos2*^−/−^ produced a much more severe phenotype than *Cybb*^−/−^. (Figure 2J–2L and Figure S2). However, the cells harboring *Mtb* in these two susceptible mouse strains differed. In wild type and *Cybb*^−/−^ mice, YFP-Mtb was evenly distributed between CD11b^+^/Ly6G^+^ neutrophils and the CD11b^+^/Ly6G-population that consists of macrophages and dendritic cells (Wolf et al., 2007). This proportion was dramatically altered in *Nos2*^−/−^ mice, where close to 90% of bacteria were found in the neutrophil compartment. (Figure 2M and 2N). Thus, unlike other susceptible mouse models, the loss of Cybb does not alter bacterial replication or the distribution of *Mtb* in different myeloid subsets. Instead, this gene plays a specific role in controlling IL1β production, neutrophil recruitment to the infected lung, and disease progression. As a result, we conclude that Cybb specifically promotes tolerance to *Mtb* infection.

### Enhanced IL-1β production by *Cybb*^−/−^ macrophages and dendritic cells is due to deregulated Caspase-1 inflammasome activation

To investigate the mechanism underlying increased IL-1β production in *Cybb*^−/−^ mice, we quantified the release of mature cytokine from bone-marrow derived macrophages (BMDMs) and bone-marrow derived dendritic cells (BMDCs). Compared to wild type, we found that *Cybb*^−/−^ BMDMs and BMDCs produced 4–5 fold more IL-1β after 24 hours of *Mtb* infection (Figure 3A and 3D). Under these conditions, wild type and *Cybb*^−/−^ cells remained equally viable and produced equivalent amounts of TNF (Figure 3B–F), suggesting that the effect of Cybb on IL-1β production was specific to this cytokine.

**Figure 3.**
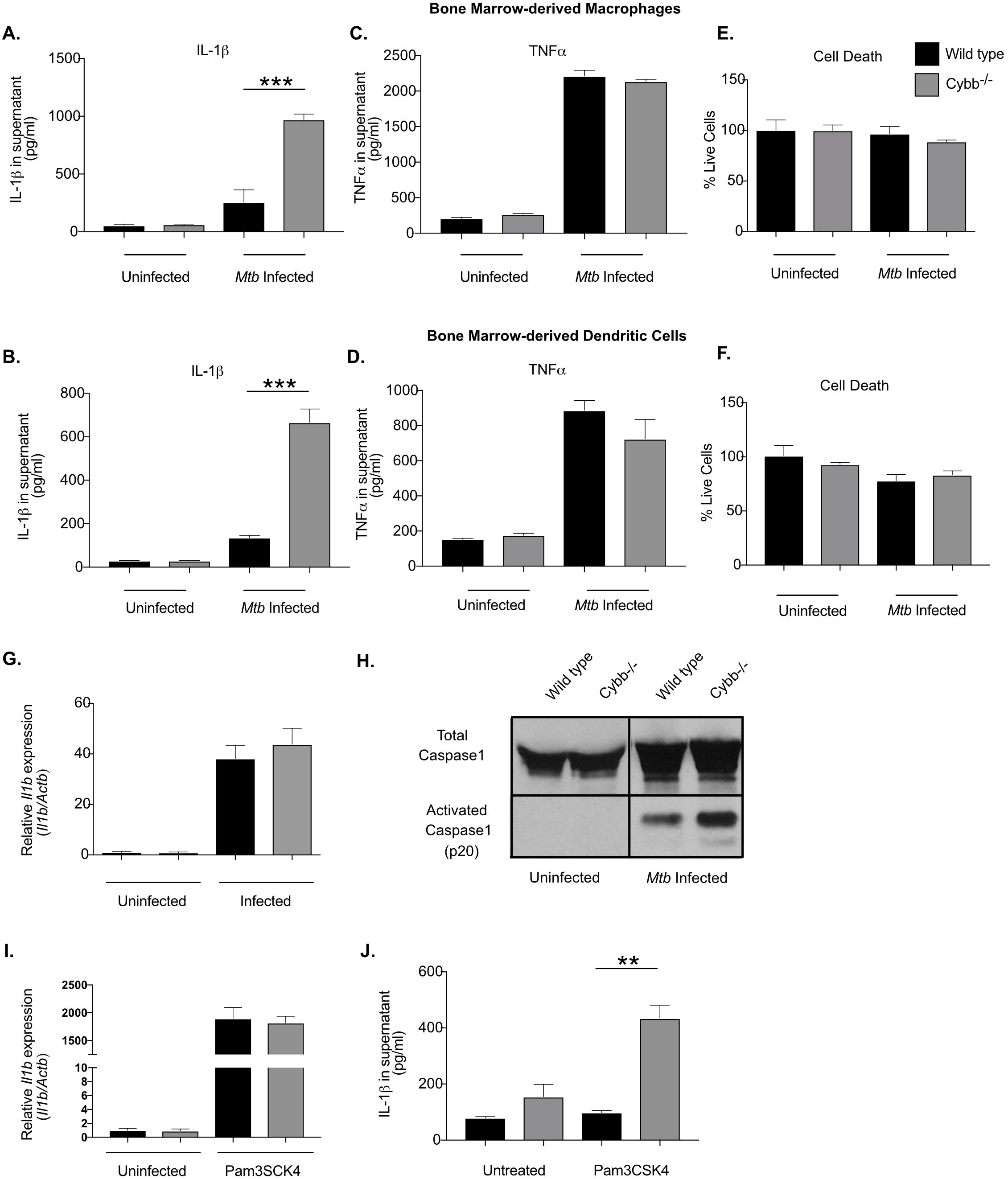
Cybb controls Caspase1 activation in macrophages and dendritic cells during *Mtb* infection. Bone marrow-derived macrophages (BMDMs) or Bone marrow-derived dendritic cells (BMDCs) from wild type or *Cybb*^−/−^ mice were left untreated or infected with *Mtb* for 4 hours then washed with fresh media. 18 hours later supernatants were harvested and the levels of **A**. and **B**. IL-1β and **C**. and **D**. TNFα were quantified in the supernatants by ELISA. Shown is the mean of 4 biological replicates normalized to a standard curve ^+/−^ s.d. *** p<.001 by unpaired two-tailed t-test. **E**. and **F**. Viability of remaining cells was determined by quantifying ATP via luminescence and compared to cells at 4 hours post-infection (mean % viability ^+/−^ s.d.). Data in A,C and E are representative of five independent experiments with at least 3 biological replicates per experiment. Data in B, D and F are representative of three independent experiments with 4 biological replicates per experiment. **G**. Relative RNA expression of *Il1b* (compared to β-Actin) was determined from wild type and *Cybb*^−/−^ BMDMs left untreated or infected for 24 hours with *Mtb* (mean ^+/−^ s.d.) by qRT-PCR. Data are representative of two independent experiments with 3–4 biological replicates per group. **H**. Immunoblot analysis was used to assess the activation of Caspase1 from Wild type and *Cybb*^−/−^ BMDMs infected for 24 hours with *Mtb*. Total Caspase1 was used as a loading control. Data are representative of 3 independent experiments with at least 3 biological replicates analyzed per experiment. **I**. Relative RNA expression of *Il1b* (compared to *Actb*) was determined from wild type and *Cybb*^−/−^ BMDMs left untreated or treated with Pam3CSK4 for 24 hours (mean ^+/−^ s.d.) by qRT-PCR. Data are representative of two independent experiments with 3–4 biological replicates per group. **J**. Wild type and *Cybb*^−/−^ BMDMs were left untreated or treated with PAM3CSK4 for 12 hours, supernatants were harvested and the levels of IL-1β were quantified by ELISA (mean ^+/−^ s.d.). ** p<.01 by unpaired two-tailed t-test. Data are representative of three independent experiments with 4 biological replicates per experiment.

The production of mature IL-1β requires two distinct signals (von Moltke et al., 2013). The first signal induces the expression of *Il1b* mRNA and subsequent production of pro-IL-1β, and a second signal activates Caspase1, which is necessary for the processing and secretion of mature IL-1β. To understand what step of IL1β production was altered in *Cybb*^−/−^ cells, we quantified these two signals. The expression of *Il1b* mRNA in uninfected and infected BMDMs was unchanged between wild type and *Cybb*^−/−^ BMDMs (Figure 3G). In contrast, under the same conditions, the processing of Caspase1 to its active form was increased in infected *Cybb*^−/−^ BMDMs compared to wild type cells (Figure 3H).

These observations suggested that caspase-1 activity is increased in *Cybb*^−/−^ cells, which could allow mature IL-1β secretion in the absence of an inflammasome activator. To test this hypothesis, we stimulated cells with the TLR2 agonist, PAM3CSK4, to induce pro-IL-1β production. PAM3CSK4 stimulation induced *Il1b* mRNA to similar levels between wild type and *Cybb*^−/−^ BMDMs, albeit over 100 times higher than infection with *Mtb* (Figure 3I). In wild type cells, this induction of *Il1b* expression produced little mature IL-1β secretion, consistent with the need for subsequent inflammasome activation. In contrast, induction of *Il1b* expression led to robust secretion of mature IL-1β from *Cybb*^−/−^ BMDM, consistent with unregulated inflammasome activity in these cells (Figure 3J). Together these data show that loss of Cybb leads to hyper-activation of Caspase1 and increased release of IL-1β during *Mtb* infection of both BMDMs and BMDCs.

### Loss of tolerance is reversed in *Cybb*^−/−^ macrophages and mice by blocking the production or activity of IL-1β

The NLRP3 inflammasome consists of NLRP3, ASC, and Caspase 1. While this complex is generally responsible for IL-1β processing in *Mtb* infected macrophages (Coll et al., 2015; Dorhoi et al., 2012), it remained unclear whether the enhanced IL-1β secretion from *Cybb*^−/−^ cells also relied on these components. To identify the responsible complex, we blocked the activation of the NLRP3 inflammasome in several distinct ways. The NLRP3 inflammasome is specifically inhibited by IFNγ stimulation, via the nitric oxide-dependent nitrosylation of the NLRP3 protein (Mishra et al., 2013). Pretreatment of wild type and *Cybb*^−/−^ BMDMs with varying concentrations of IFNγ, inhibited the secretion of IL-1β from both wild type and *Cybb*^−/−^ BMDMs compared to untreated cells. While this result indicated an important role for NLRP3, IFNγ pretreatment did not completely suppress IL-1β secretion and there remained significant differences in the IL-1β release between *Cybb*^−/−^ and wild type cells at all concentrations of the cytokine.

To more directly assess the role of NLRP3 and Caspase-1 in IL-1β production by *Cybb*^−/−^ cells, we employed specific small molecule inhibitors. Treatment of Mtb-infected BMDMs with either the NLRP3 inhibitor, MCC950 (Coll et al., 2015), or the Caspase-1 inhibitor, VX-765 (Stack et al., 2005) caused a dramatic reduction in IL-1β in both wild type and *Cybb*^−/−^ BMDMs compared to untreated cells (Figure 4B and 4C). This ten-fold decrease in IL-1β secretion could not be attributed to inhibition of pro-IL-1β levels, as none of these inhibitors affected *Il1b* mRNA by more than two-fold. Similarly, the spontaneous IL-1β secretion observed upon PAM3CSK4 stimulation was also inhibited by MCC950 and IFNγ, (Figure 4E). In each case inflammasome inhibition reduced IL-1β secretion to the same level in both wild type and *Cybb*^−/−^ cells, indicating that the NLRP3 inflammasome was responsible for the enhanced production of this cytokine in *Cybb*^−/−^ BMDM.

**Figure 4.**
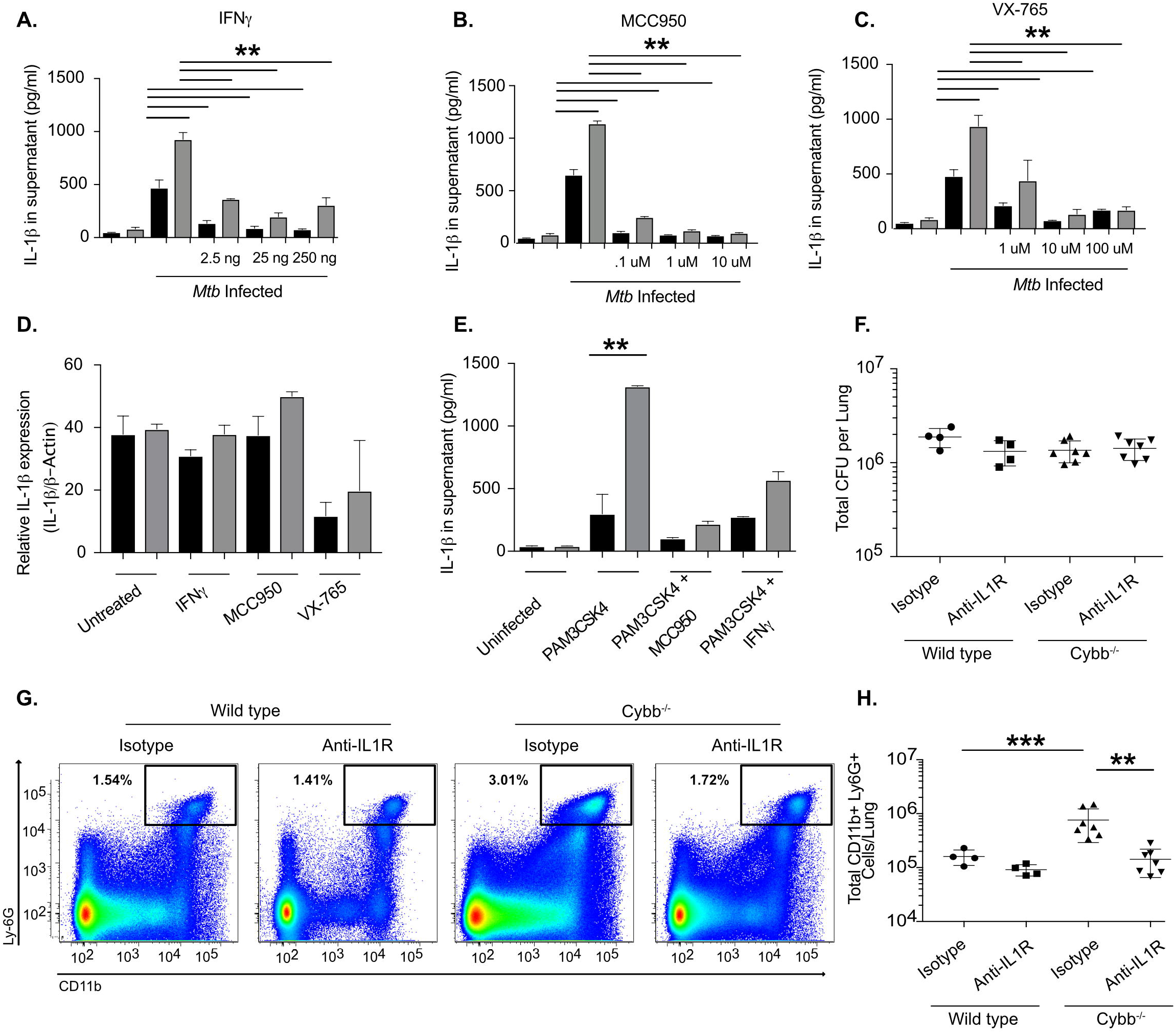
Hyper-inflammation in *Cybb*^−/−^ is reversed by inflammasome and IL1 inhibition. **A**. Wild type (Black Bars) and *Cybb*^−/−^ (Grey Bars) BMDMs were left untreated or treated with with the indicated concentrations of IFNγ for 12 hours. Cells were then infected with *Mtb* for 4 hours then washed with fresh media. 18 hours later supernatants were harvested and levels of IL-1β from each condition were quantified by ELISA (mean ^+/−^ s.d.). ** p-value <.01 * p-value <.05 by one-way ANOVA with tukey correction. Data are representative of three independent experiments with at least 3 biological replicates per experiment. **B**. Wild type and *Cybb*^−/−^ BMDMs were left untreated or treated with the indicated concentrations of MCC950 for 2 hours. Cells were then infected with *Mtb* for 4 hours then washed with fresh media with inhibitor. 18 hours later supernatants were harvested and levels of IL-1β from each condition were quantified by ELISA (mean ^+/−^ s.d.). ** p-value <.01 by one-way ANOVA with tukey correction. Data are representative of three independent experiments with at least 3 biological replicates per experiment. **C**. Wild type and *Cybb*^−/−^ BMDMs were left untreated or treated with the indicated concentrations of VX-765 for 2 hours. Cells were then infected with *Mtb* for 4 hours then washed with fresh media with inhibitor. 18 hours later supernatants were harvested and levels of IL-1β from each condition were quantified by ELISA (mean ^+/−^ s.d.). ** p-value <.01 by one-way ANOVA with tukey correction. Data are representative of three independent experiments with at least 3 biological replicates per experiment. **D**. Relative RNA expression of IL-1β (compared to b-Actin) was determined from wild type and *Cybb*^−/−^ BMDMs left infected for 24 hours with *Mtb* in the presence or absence of the indicated inhibitors (mean ^+/−^ s.d.) by qRT-PCR. **E**. Wild type and *Cybb*^−/−^ BMDMs were left untreated or treated 25ng/ml IFNγ or 1μM MCC950 overnight. The following day cells were treated with PAM3CSK4 for 12 hours, supernatants were harvested and the levels of IL-1β were quantified by ELISA (mean ^+/−^ s.d.). ** p<.01 by unpaired two-tailed t-test. Data are representative of two independent experiments with 4 biological replicates per experiment. **F**. Wild type and *Cybb*^−/−^ mice were infected intratracheally with *Mtb* strain 18b and treated for two weeks daily with streptomycin then were injected every other day for two weeks with 200ug of either isotype control antibody or anti-IL1R antibody. The total levels of viable *Mtb* in the lungs was determined by CFU plating on streptomycin (mean ^+/−^ s.d.). **G**. Representative flow cytometry plot showing Ly6G^+^ CD11b^+^ neutrophil recruitment to the lungs of *Cybb*^−/−^ mice during control and IL1R blockade conditions (gated on live/singlets/CD45^+^). **H**. Quantification of neutrophil recruitment to the lungs at the indicated time points following infection for wild type and *Cybb*^−/−^ mice during control and IL1R blockade conditions is shown as an absolute number of Ly6G^+^ CD11b^+^ cells per lung (mean ^+/−^ s.d). *** p-value <.001 ** p-value <.01 by one-way ANOVA with tukey correction. Data in F-G are representative of two independent experiments with 4–7 mice per group.

Based on these studies, we hypothesized that the tolerance defect observed in the intact mouse was due to inflammasome-dependent IL-1 signaling. IL-1β serves a complex role during infection (Mayer-Barber et al., 2010; Nunes-Alves et al., 2014). Some production of this cytokine is important for the antimicrobial immunity, but persistent production can lead to pathology. In order to focus on the role of over-production of IL-1β on tolerance, we inhibited IL-1 signaling in mice infected with non-replicating auxotrophic *Mtb* to normalize the bacterial burden. Two weeks after infection, wild type and *Cybb*^−/−^ mice were treated with either an isotype control antibody or an anti-IL1R antibody to block the effect of increased IL-1β production. As expected, *Mtb* levels were similar in all mice, but more neutrophils accumulated in the lungs *Cybb*^−/−^ animals (Figure 4F–H). While anti-IL-1 R treatment had no effect in wild type animals, inhibition of IL-1 signaling reduced neutrophil infiltration in *Cybb*^−/−^ mice to the level observed in wild type animals. Taken together, our data show that *Cybb*^−/−^ contributes to protective immunity to *Mtb* not by controlling bacterial replication, but instead by preventing an IL1 dependent inflammatory response that increases neutrophil recruitment to the lung and exacerbates disease progression.

## Discussion

The role of the Phox complex in protection from TB has presented a paradox (Deffert et al., 2014a). Based on the well-described antimicrobial properties of Phox-derived ROS, previous studies have focused on examining the function of Phox components in controlling *Mtb* replication in mice (Cooper et al., 2000; Deffert et al., 2014b; Jung et al., 2002). The minimal effects observed in these assays suggested that Phox may not play a protective role in TB. This conclusion contrasts with several studies indicating that human CGD patients show increased susceptibility to TB disease (Bustamante et al., 2011; Deffert et al., 2014a; Khan et al., 2016). Our dissection of disease progression in Cybb-deficient mice harmonizes these conflicting observations. While we verify that Phox plays no discernable role in antimicrobial resistance to *Mtb*, we uncovered a previously unknown role for this complex in promoting tolerance to *Mtb* infection and inhibiting TB disease progression.

While we were able to clearly delineate the role of Phox during *Mtb* infection, the role(s) played by this complex in any given infection is likely to vary. Phox-deficient mice are unable to control the growth of several bacterial pathogens that are known to cause serious infections in CGD patients, including non-tuberculous mycobacteria (Deffert et al., 2014b; Dinauer et al., 1997; Fujita et al., 2010; Jackson et al., 1995). In the context of these infections, the antimicrobial functions of Phox may predominate. In contrast, for pathogens such as *Mtb* that are resistant to ROS-mediated toxicity and persist in the tissue to promote continual inflammatory damage, the tolerance-promoting activity of Phox appears to play a dominant role.

During *Mtb* infection, we found that the ROS produced by Phox are critical to control the activation of the NLRP3 inflammasome. In contrast, mitochondrial ROS are well known to activate inflammatory cascades (Weinberg et al., 2015), suggesting that the context by which ROS are produced influences the inflammatory outcome of activated cells. Despite this complexity, Phox-deficient mice and CGD patients suffer from hyper-inflammatory diseases including arthritis, colitis, and prolonged inflammatory reactions to microbial products, indicating that the dominant immunoregulatory role for Phox-derived ROS is anti-inflammatory (Morgenstern et al., 1997; Schappi et al., 2008; Segal et al., 2010). Several non-mutually exclusive mechanisms could explain this anti-inflammatory effect. For example, the loss of Phox derived ROS has been proposed to promote the production of inflammatory mediators by inhibiting autophagy (de Luca et al., 2014). Another mechanism was described in superoxide dismutase1 (*Sod1*) deficient cells, where the accumulation of ROS inhibits Caspase1 activation through glutathionation of reactive cysteines (Meissner et al., 2008). This latter mechanism is reminiscent of the process by which nitric oxide (NO), inhibits inflammasome activation via S-nitrosylation of NLRP3 (Mishra et al., 2013). The intersection of these two important anti-inflammatory pathways at the NLRP3 inflammasome indicates that this complex may be a critical point of integration where inflammatory cascades are controlled during chronic infections.

Our findings are consistent with a growing body of literature suggesting that inflammasome-derived IL-1 promotes TB disease progression. For example, genetic polymorphisms that increase the expression of IL1β or the production of IL-1 dependent pro-inflammatory lipid mediators are associated with TB disease progression (Mishra et al., 2017; Zhang et al., 2014). Similarly, transcriptional signatures of inflammasome activation have been observed in severe forms of TB disease, such as meningitis (Marais et al., 2017) and TB-associated immune reconstitution syndrome (Tan et al., 2016). Together with our work, these findings imply an important role for infection tolerance in protection from TB disease, and implicate Caspase-1 as a critical point at which tolerance is regulated.

## Materials and Methods

### Mice

C57BL/6J (Stock # 000664), *Cybb*^−/−^ (B6.129S-Cybb^tm1Din^/J stock # 002365), Nos2^−/−^ (B6.129P2-Nos2^tm1Lau^/j, stock # 002609), B6.SJL-*Ptprc^a^ Pepc^b^* carrying the pan leukocyte marker CD45.1 (stock # 002014) were purchased from the Jackson Laboratory. Mice were housed under specific pathogen-free conditions and in accordance with the University of Massachusetts Medical School, IACUC guidelines. All animals used for experiments were 6–12 weeks except mixed chimeras that were infected at 16 weeks following 8 weeks of reconstitution.

### Mouse Infection

Wild type M. tuberculosis strain H37Rv was used for all studies unless indicated. This strain was confirmed to be PDIM-positive. Prior to infection bacteria were cultured in 7H9 medium containing 10% oleic albumin dextrose catalase growth supplement (OADC) enrichment (Becton Dickinson) and 0.05% Tween 80. H37Rv expressing msfYFP has been previously described and the episomal plasmid was maintained with selection in Hygromycin B (50ug/ml) added to the media (Mishra et al., 2017). For low and high dose aerosol infections, bacteria were resuspended in phosphate-buffered saline containing tween 80 (PBS-T). Prior to infection bacteria were sonicated then delivered via the respiratory route using an aerosol generation device (Glas-Col). Infections of mice with the streptomycin auxotrophic strain of *Mtb* (18b) have been previously described. In short mice were infected via intra-tracheal infection and treated daily with 2mg of streptomycin daily for two weeks. For anti-IL1R treatment mice were injected with 200ug of anti-IL1R antibody or Isotype control (Bio-xcell) every other day starting at day 14. Both male and female mice were used throughout the study and no significant differences in phenotypes were observed between sexes.

### Immunohistochemistry

Lungs from indicated mice were inflated with 10% buffered formalin and fixed for at least 24 hours then embedded in paraffin. Five-Micrometer—thick sections were stained with haematoxylin and eosin (H&E). All staining was done by the Diabetes and Endocrinology Research Center Morphology Core at the University of Massachusetts Medical School.

### Flow Cytometry

Lung tissue was harvested in DMEM containing FBS and placed in C-tubes (Miltenyi). Collagenase type IV/DNaseI was added and tissues were dissociated for 10 seconds on a GentleMACS system (Miltenyi). Tissues were incubated for 30 minutes at 37C with oscillations and then dissociated for an additional 30 seconds on a GentleMACS. Lung homogenates were passaged through a 70-micron filter or saved for subsequent analysis. Cell suspensions were washed in DMEM, passed through a 40-micron filter and aliquoted into 96 well plates for flow cytometry staining. Non-specific antibody binding was first blocked using Fc-Block. Cells were then stained with anti-Ly-6G Pacific Blue, anti-CD4 Pacific Blue, anti-CD11b PE, anti-CD11c APC, anti-Ly-6C APC-Cy7, anti-CD45.2 PercP Cy5.5, anti-CD3 FITC, anti-CD8 APC-Cy7, anti-B220 PE-Cy7 (Biolegend). Live cells were identified using fixable live dead aqua (Life Technologies). For infections with fluorescent H37Rv, lung tissue was prepared as above but no antibodies were used in the FITC channel. All of these experiments contained a non-fluorescent H37Rv infection control to identify infected cells. Cells were stained for 30 minutes at room temperature and fixed in 1% Paraformaldehyde for 60 minutes. All flow cytometry was run on a MACSQuant Analyzer 10 (Miltenyi) and was analyzed using FlowJo_V9 (Tree Star).

### Macrophage and Dendritic Cell Generation

To generate bone marrow derived macrophages (BMDMs), marrow was isolated from femurs and tibia of age and sex matched mice as previously described. Cells were then incubated in DMEM (Sigma) containing 10% fetal bovine serum (FBS) and 20% L929 supernatant. Three days later media was exchanged with fresh media and seven days post-isolation cells were lifted with PBS-EDTA and seeded in DMEM containing 10% FBS for experiments.

To generate bone marrow derived dendritic cells (BMDCs) Marrow was isolated from femurs and tibia of age and sex matched mice. Cell were then incubated in iMDM media (GIBCO) containing 10% FBS, L-Glutamine, 2μM 2-mercaptoethanol and 10% B16-GM-CSF supernatant (Zanoni et al., 2016). BMDCs were then purified on day six using Miltenyi LS columns first using negative selection for F480 followed by CD11c positive selection. Cells were then plated and infected the following day.

### Macrophage and Dendritic Cell Infection

*Mtb* or *Mycobacterium bovis-BCG* were cultured in 7H9 medium containing 10% oleic albumin dextrose catalase growth supplement (OADC) enrichment (Becton Dickinson) and 0.05% Tween 80. Before infection cultures were washed in PBS-T, resuspended in DMEM containing 10%FBS and passed through a 5-micron filter to ensure single cells. Multiplicity of infection (MOI) was determined by optical density (OD) with an OD of 1 being equivalent to 3×10^8^ bacteria per milliliter. Bacteria were added to macrophages for 4 hours then cells were washed with PBS and fresh media was added. At the indicated time points supernatants were harvested for cytokine analysis and the cells were processed for further analysis. Cell death was assessed using Cell-Titer-Glo luminescent cell viability assay (Promega) following manufacturer’s instructions. For inhibitor treatments cells were treated with the indicated concentrations of IFNγ (Peprotech), MCC950 (Adipogen) or VX-765 (Invivogen) or vehicle control overnight prior to infection and maintained in the media throughout the experiment.

### Mixed Bone Marrow Chimera generation and cell sorting

Mixed bone marrow chimera experiments were done essentially as previously described. Wild-type CD45.1^+^ mice were lethally irradiated with two doses of 600 rads. The following day, bone marrow from CD45.1^+^ wild-type mice and CD45.2^+^ knockout mice (wild-type or Cybb^−/−^) was isolated, red blood cells were lysed using Tris-buffered ammonium chloride (ACT), and the remaining cells were quantified using a haemocytometer. CD45.1^+^ and CD45.2^+^ cells were then mixed equally at a 1:1 ratio and 10^7^ cells from this mixture were injected intravenously into lethally irradiated hosts that were placed on sulfatrim for three weeks. 8 weeks later mice were then infected by low-dose aerosol with *M. tuberculosis* H37Rv. Four weeks following infection, the lungs of chimera mice were processed for flow cytometry. An aliquot of this suspension was saved for flow cytometry analysis of the lung population and overall bacterial levels. The remaining cells were split equally and stained with either anti-CD45.1 APC or anti-CD45.2 PE.

Stained populations were then incubated with either anti-APC or anti-PE magnetic beads (Miltenyi) following the manufacturer’s instructions and sorted using LS-columns (Miltenyi). Purified cells were divided equally and then plated for M. tuberculosis on 7H10 agar or counted and stained for analysis of cellular purity. Cells from the input homogenate, flow through and the positive sort fractions were stained with for purity. Samples with >90% were used for subsequent analysis. At 21 days after plating, colonies were enumerated and the *Mtb* levels per sorted cells were determined.

### qRT-PCR

Cells were lysed in Trizol-LS (Thermofisher), RNA was purified using Direct-zol RNA isolation kits (Zymogen) and quantified on nanodrop. RNA was diluted to 5ng/μl and 25 ng total RNA was used for each reaction. Ct values for each sample were determined in technical duplicates for β-Actin and IL-1β using one-step RT-PCR Kit (Qiagen) on a Viia7 Real-time PCR system (Life Technologies). ΔΔct values were then determined for each sample.

### Immunoblotting and Cytokine quantification

Murine cytokine concentrations in culture supernatants and cell-free lung homogenates were quantified using commercial enzyme-linked immunosorbent assay (ELISA) kits (R&D). All samples were normalized for total protein content. Caspase1 activation in macrophage lysates was determined by western blotting with Caspase1 antibody purchased from Adipogen.

## Author Contributions

AO and CMSassetti conceived of and designed the experiments. AO, CMSmith, and MK performed the experiments and analyzed the data. AO and CMSasssetti wrote the initial manuscript. All authors edited the manuscript.

## Acknowledgements

We are thankful to members of the Sassetti and Behar lab for helpful discussions. This work was funded by the Arnold and Mabel Beckman Postdoctoral Fellowship (AO), Charles King Trust Postdoctoral Fellowship (CMSmith), and NIH Grant AI132130 (CMSassetti)

---

**Figure S1. A**. Gating strategy used for all flow cytometry analysis. Shown is a representative gating approach for the analysis used throughout the paper. Cells were first gated on live/dead negative cells then forward by side scatter then singlets. We then gated on all CD45+ cells in and analyzed the subsequent populations using the indicated fluorescent markers. **B**. Quantification of macrophages, B cells, Dendritic cells and T cells recruitment to the lungs at the 28 days following infection for wild type and *Cybb*^−/−^ mice is shown as absolute number of cells per lung (mean ^+/−^ s.d).

**Figure S2. A**. A representative sample of the purity of the cell populations depicted in “Figure 2b and c” were determined in the input and sorted fraction of MACS purification of the lung leukocytes from mixed-bone marrow chimeric mice. **B**. Lung homogenates from wild type, *Cybb*^−/−^ or *Nos2*^−/−^ mice infected with *Mtb* for 28 days were probed for the cytokines IL-1β by ELISA (mean ^+/−^ s.d). Results are representative of 3 independent experiments with 3–5 mice per group. ** p-value <.01 by unpaired two-tailed t-test.

